# Managing the genetic diversity of animal populations using cryobanks: optimizing the constitution of *ex situ* collection?

**DOI:** 10.1101/2025.02.19.638995

**Authors:** Alicia Jacques, Michèle Tixier-Boichard, Gwendal Restoux

**Affiliations:** Université Paris-Saclay, INRAE, AgroParisTech, GABI, 78350 – Jouy-en-Josas, France; Eliance, 149 rue de Bercy, 75012 – Paris, France

**Keywords:** gene bank, simulations, biodiversity, breeding scheme, genetic resources

## Abstract

To compensate for the erosion of genetic variability taking place in local breeds as well as in large breeds under selection, many countries have set up cryopreserved collections. Currently, there are as many sampling strategies as there are cryobanks. The aim of our study was to examine some parameters for optimizing the constitution of a gene bank.

**Results:** We simulated different breeding scenarios of animal populations (*i.e*. selection or conservation) and examined the impact of using cryopreserved collections according to (i) the size and age of collections or (ii) the sampling strategy to set up these collections.

The age of cryopreserved resources was the predominant factor of use in populations under selection, where the use of recently collected sires was favored. For populations under conservation, older reproductive material becomes highly relevant. Using a large number of old individuals enabled better management of matings and helped to slow down the increase in inbreeding on the long term, while the use of a lower number of old individuals enabled some genetic progress to be maintained. Since a gene bank needs to address the needs of all types of populations, continuous enrichment of collections appeared to be a relevant option. In addition, the use of OCS can help to efficiently mobilize *ex situ* genetic resources in animal breeding programs.

**Conclusions:** Cryopreserved genetic resources are an invaluable help in the management of domestic populations. Relevant recommendations for the management of cryopreserved collections can be made according to breeding programs.

## Introduction

Genetic diversity is an essential component in the evolution and adaptation of populations. Many studies have investigated the impact of a loss in genetic variability for populations at different time scales (Frankham, 2005; Wiggans et al., 2017; Kardos et al., 2021). In the long term, allelic variability among individuals favors the adaptive potential of a population (Notter, 1999; Reed and Frankham, 2003; de Cara et al., 2013) by increasing the probability that some genotypes will be advantageous in face of a disturbance (*e.g*. emergence of a new pathogen, new environmental constraints *etc*…). In the medium term, the erosion of genetic diversity has an impact on selection response, in particular on genetic progress for domestic populations (Kinghorn et al., 2009; Meuwissen et al., 2013; Woolliams and Oldenbroek, 2017). In the short term, reduced variability leads to inbreeding depression, *i.e*. a reduction of selective value (O’Grady et al., 2006; Charlesworth and Willis, 2009; Leroy, 2014). Consequently, whatever the type of population, conserving genetic diversity is needed.

For several decades, initiatives have been taken to conserve genetic diversity of domestic animal breeds across countries. This can be done by a careful management of on-farm populations, also called ‘*in situ*’ conservation, and/or through the development of gene banks,using cryopreservation of reproductive material, also called ‘*ex situ’* conservation. The Food and Agriculture Organization of the United Nations (FAO) has identified cryoconservation of animal genetic resources as an essential element in national conservation strategies (FAO, 2007; FAO, 2015). Over the last decades, germplasm collections have been developed for many breeds of farm animals in different countries (Blackburn, 2018; Leroy et al., 2019; Jacques et al., 2024). In 2016, the IMAGE H2020 project (Innovative Management of Animal Genetic Ressources) developed new methods, such as the MoBPS simulation tool, to support the use of genetic collections and the management of animal genebanks. In the same way, the CRB-Anim project (French Biological Animal Resource Center, https://doi.org/10.15454/1.5613785622827378E12) has enabled the enrichment of cryopreserved collections for various animal species and breeds. These initiatives have developed methodological and technical skills to enhance germplasm collections, and have provided financial support for collection enrichment in some instances. Yet, a major question remains: how to set-up a gene bank, for which purposes?

Gene banks can help restore breeds in the event of extinction, provided that sufficient genetic material has been preserved, as described by Windig *et al*. (2025) with the proposal of “the conservation planner” in order to assess the number of samples to be stored in a gene bank to enable the reconstruction of a breed. Extinction of a breed is the worst case. Both ‘*in situ*’ and ‘*ex situ’* approaches are complementary and the use of gene bank collections can help manage genetic variability and genetic progress in selected populations (Leroy et al., 2011; Doekes et al., 2018; Eynard et al., 2018; Jacques et al., 2026). A few real cases of using cryopreserved semen in breeding schemes have been described for two local pig breeds in France (Mercat et al., 2008), for reintroducing genetic diversity of the Y chromosome in Holstein population in the United States (Dechow et al., 2020) and for a local French dairy cattle breed (Jacques et al., 2023).

Thus, gene banks can be used for a range of purposes which should be kept in mind when collecting reproductive material for long-term preservation. Whereas FAO has proposed recommendations on the genetic material required to reconstitute a breed (FAO, 2012), there are no such recommendation for other uses, such as reintroducing lost diversity in an existing population. Furthermore, long term cryopreservation of animal genetic resources has a cost (Pizzi et al., 2016; De Oliveira Silva et al., 2019). Strategies are needed to help gene banks anticipating future uses of their collections while considering practical issues such as the number of samples and the timing of collection enrichment.

To our knowledge, no study has investigated the impact of different gene banking strategies for animal genetic resources according to breeding programs. In this study, we are using simulations to compare the use of gene banks collections which were assembled in different ways.

## Methods

We performed stochastic simulations of different breeding schemes using the R package MoBPS, version 1.6.64 (Pook et al., 2020; Pook, 2021). R version 4.2.1 (R Core team, 2020) and ggplot2 R packages (Wickham, 2011) were used to perform all additional calculations and graphical representations. Statistical tests were performed using the lm function, post-hoc tests were conducted using the emmeans package (Lenth et al., 2021), and type II ANOVA were performed using the car package (Fox et al., 2019).

### Definition of scenarios: type of selection, prolificity and uses of germplasm collections

We used the same four scenarios as in Jacques *et al*., (2026), considering two traits of interest (Trait 1 and Trait 2) both having a heritability of 0.4 and exhibiting a negative genetic correlation of −0.3.

The scenario rm corresponds to a random choice of parents among the candidates; the scenario max_BV corresponds to a choice of parents that maximizes genetic gain on a synthetic index combining both traits; the scenario max_GD corresponds to a choice of parents that minimizes the loss of genetic diversity; and the scenario OCS corresponds to a choice of parents that maximizes genetic gain under the constraint of a maximal increase in kinship of 0.5% per generation, as recommended by the FAO (FAO, 1998). The max_BV, max_GD, and OCS scenarios were modeled using the optimal contribution strategy (OCS) (Meuwissen, 1997) implemented in MoBPS via the optiSel package, version 2.0.5 (Wellmann, 2019; Wellmann, 2021). The relationship matrix in the OCS was calculated using the vanRaden method (VanRaden, 2008).

The four scenarios were simulated as described in Jacques *et al*., (2026), with two options: using the contemporary candidates only, or adding 40 cryopreserved sires to the pool of candidates. In the latter case, cryopreserved individuals were systematically re-evaluated at the same time as contemporary individuals. All scenarios were replicated 20 times.

### Initialization of simulated populations and pre-burn-in

The pre-burn-in and burn-in steps were performed as described in Jacques *et al*., (2026). The size of the breeding scheme was also the same, namely a founder population of 500 individuals with a sex ratio of 0.5, with 5000 genetic markers (SNPs) regularly distributed along the genome.

#### Genetic architecture and definition of a synthetic index

Each trait was simulated as in Jacques *et al*., (2026) with 500 additive quantitative trait loci (QTLs). The estimated genetic values of individuals (EBVs) were obtained using GBLUP evaluation models incremented in MoBPS software. The true breeding values (BVs) were generated by the genotype simulation and used to standardize the EBV for each trait. Then, EBVs were combined in a synthetic index using a weight of 0.8 for Trait 1 and a weight of 0.2 for Trait 2. Thus, for each sire, the index value was obtained as follows:

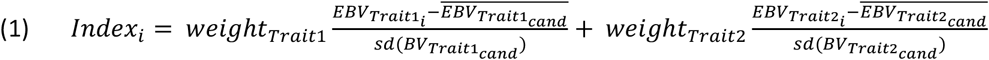

Where *index*_*i*_ is the synthetic index value of animal *i, weight*_*Trait*1_ and *weight*_*Trait*2_ are the weights of Trait1 and Trait2 in the selection objective, respectively (*i.e*. here 0.8 and 0.2).

#### Pre-burn-in and initial population creation

The pre-burn-in step generated linkage disequilibrium and genetic structure for 20 simulated populations during 10 generations (Figure 1). Each new cohort included 250 males and 250 females resulting from the mating of the 20 best males and the 250 best females of the previous cohort, with prolificacy set at 2, half-sib matings prohibited, and the selection of sires based on the synthetic index defined above.

**Figure 1.**
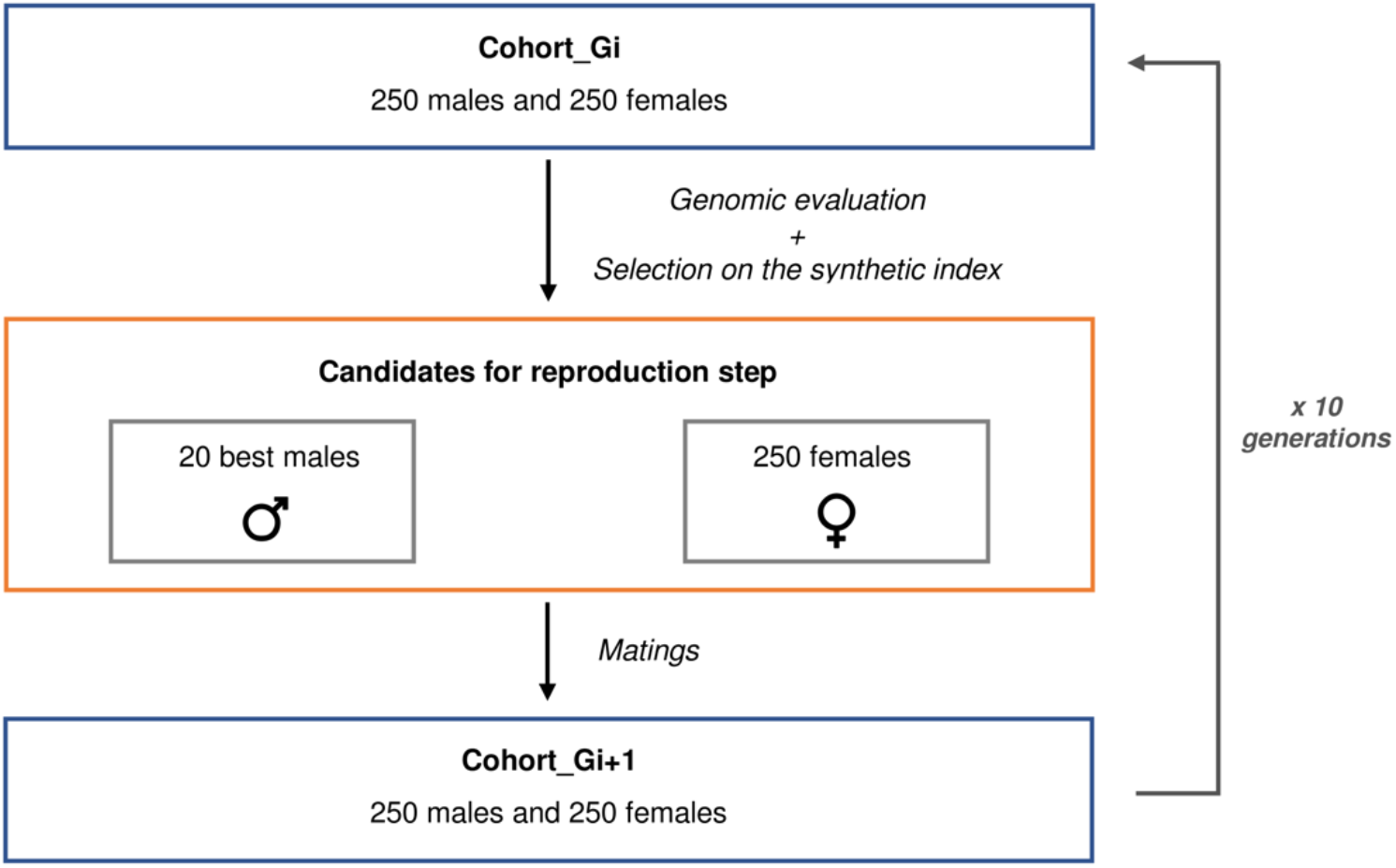
Breeding steps for the pre-burn-in of simulations.

### Burn-in phase and breeding program

The burn-in step was replicated 20 times over 10 overlapping generations, for two breeding programs: population under selection or population under conservation(Figure 2). In the case of selection, candidate sires were selected on the basis of the synthetic index, as defined above in equation (1).

**Figure 2.**
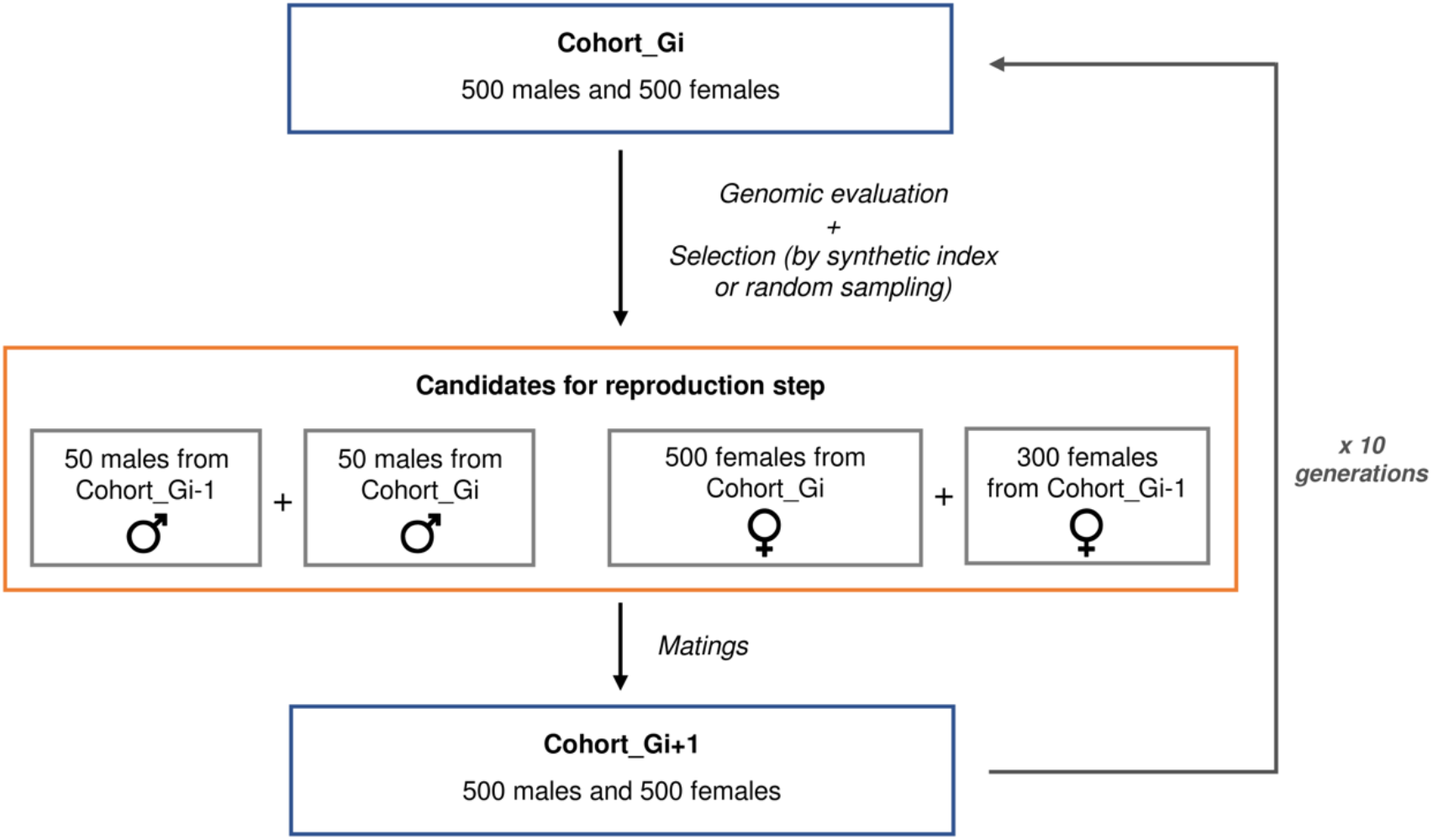
Breeding steps for the selection or conservation burn-in.

The target was to obtain an average kinship in the 20^th^ generation of (i) 14–15% in the case of a population under selection or (ii) 10–11% in the case of a population under conservation.

### Definition of germplasm collection types

#### Discontinous sampling with variable collection size also names as ‘fixed cryobanks’

Individuals were randomly selected among sires of a given generation to create the cryopreserved collections.

Three types of collections were defined according to the age of the cryopreserved sires: (a) an ancient cryobank composed of individuals cryopreserved from early generations 10, 11, 12 and 13, (b) a recent cryobank composed of individuals cryopreserved from late generations 15, 16, 17 and 18 and (c) a mixed cryobank composed of individuals cryopreserved across generations 11, 13, 15 and 17. The aim was to study whether the age of the cryopreserved sires had an impact on their use.

Then, we varied the number of individuals cryopreserved in each generation, storing 5, 10 or 20 individuals, so that the size of collections varied from 20, to 40 or 80 cryopreserved sires. The aim was to study whether the size of the collection had an impact on the use of *ex situ* genetic resources.

#### Continous sampling, also named as ‘mobile cryobanks’

Three types of collections were simulated.

The first two, either recent of continuous, had a fixed size with 40 individuals sampled for a given generation, noted *gen*, but a variable composition over time, as they were regularly updated, with older individuals being eliminated in favor of more recent ones joining the collections. This mimics the dynamic management under the constraint of a fixed-budget or a fixed storage capacity. The recent mobile cryobank was composed of ten cryopreserved individuals from the last four generations that were no longer proposed as sires to create the *gen*+1 cohort: *gen*-2, *gen*-3, *gen*-4 and *gen*-5. The continuous mobile cryobank was composed of five cryopreserved individuals from the last eight generations which were no longer proposed as sires to create the *gen*+1 cohort: *gen*-2, *gen*-3, *gen*-4, *gen*-5, *gen*-6, *gen*-7, *gen*-8 and *gen*-9. An example of these two types is shown in Figure 3.

**Figure 3.**
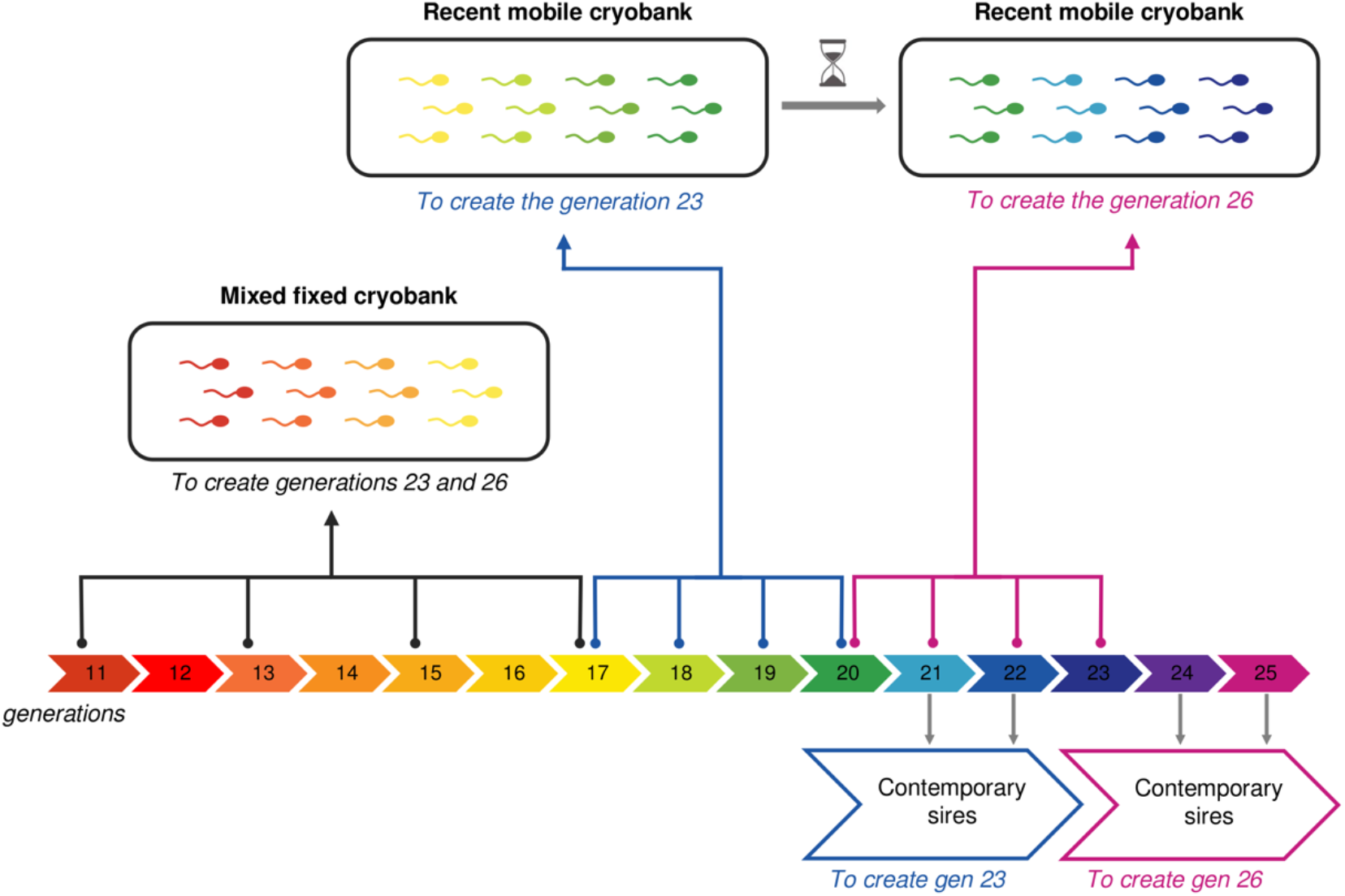
Breeding steps for the selection or conservation burn-in.

The third type is the cumulative mobile cryobank, the size of which varies from generation to generation, by adding the ten individuals from *gen*-2 to the collection at each generation. In this way, the cryobank was enriched every generation, while the other two types of mobile cryobank operated at constant size.

The use of cryobank collections was then studied for each breeding program and scenario, with either selection or conservation. Results were analysed to identify which types of collections best met the expectations of the different scenarios.

### Criteria for comparing scenarios

#### Use of cryopreserved individuals

We quantified the effective use of cryopreserved individuals by counting their average number of direct descendants (i.e., sons or daughters, i.e. rank 1), as well as indirect descendants (e.g., grandchildren, i.e. rank 2, great-grandchildren, i.e. rank 3). Numbers of offsprings resulting from the use of cryopreserved sires were analysed according to the number of generations (i.e; the genealogical rank) separating a sire from its offsprings, and compared across scenarios.

#### Breeding values

EBVs for traits 1 and 2, as well as the value of the sires’ standardized synthetic index, were evaluated at each generation. We also calculated the “true” selection index, i.e., based on EBVs without standardization, for each generation by applying the respective weights of each trait in the selection objective. Trends in EBVs and the synthetic index were compared across scenarios.

#### Genetic diversity

Average kinship was calculated for each generation using the “kinship.emp.fast” function of the MoBPS package, which estimates the real kinship based on the recombination points of genotypes by sampling the population, as described in Jacques *et al*., (2026).

Levels of genetic diversity were compared from the start of the program (generation 20) to the one at the end of the program (generation 35). The genetic distance between generation 20 and generation 35 was estimated for each scenario using Nei’s method (Nei, 1972) with MoBPS’s “get.distance” function, as described in Jacques *et al*. (2026), using the following equation:

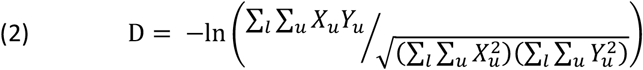

with X and Y the two population at the 20^th^ and 35^th^ generations and *X*_*u*_ (or respectively *Y*_*u*_) the frequencies for the allele *u* of the locus *l* in the population X (or respectively in the population Y).

## Results

### Population under conservation

Two scenarios were considered for this type of population: random mating (rm) and maximisation of genetic diversity (max_GD).

#### Quantifying the use of collections

Age of fixed collections had no impact on their use in conservation programs aimed purely at optimizing genetic diversity (Table 1 and Figure 4). All the proposed individuals are used, and the resulting number of offsprings is significantly different according to the number of sires used (2-factor ANOVA test, F= 628.8, df=171, p-val<0.05 for rm and F= 1188, df=171, p-val<0.05 for max_DG). The larger the collection, the greater its contribution to reproduction.

**Table 1.**
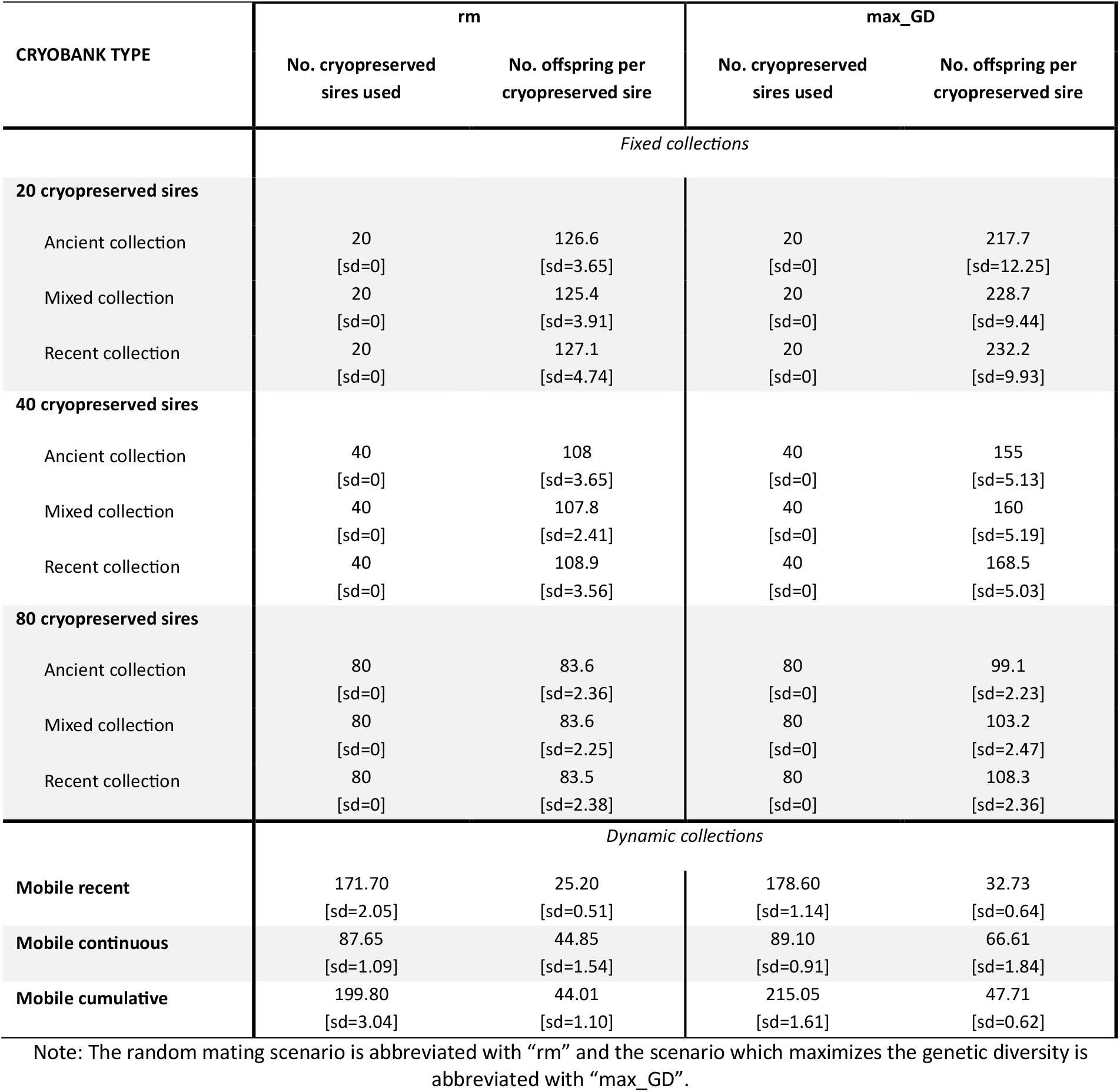
Number of cryopreserved individuals used and direct offsprings produced in populations under conservation (rm and max_GD)

| CRYOBANK TYPE | *rm* |  | *max_GD* |  |
| --- | --- | --- | --- | --- |
|  | No. cryopreserved sires used | No. offspring per cryopreserved sire | No. cryopreserved sires used | No. offspring per cryopreserved sire |
| *Fixed collections* |  |  |  |  |
| **20 cryopreserved sires** |  |  |  |  |
| Ancient collection | 20<br>[sd=0] | 126.6<br>[sd=3.65] | 20<br>[sd=0] | 217.7<br>[sd=12.25] |
| Mixed collection | 20<br>[sd=0] | 125.4<br>[sd=3.91] | 20<br>[sd=0] | 228.7<br>[sd=9.44] |
| Recent collection | 20<br>[sd=0] | 127.1<br>[sd=4.74] | 20<br>[sd=0] | 232.2<br>[sd=9.93] |
| **40 cryopreserved sires** |  |  |  |  |
| Ancient collection | 40<br>[sd=0] | 108<br>[sd=3.65] | 40<br>[sd=0] | 155<br>[sd=5.13] |
| Mixed collection | 40<br>[sd=0] | 107.8<br>[sd=2.41] | 40<br>[sd=0] | 160<br>[sd=5.19] |
| Recent collection | 40<br>[sd=0] | 108.9<br>[sd=3.56] | 40<br>[sd=0] | 168.5<br>[sd=5.03] |
| **80 cryopreserved sires** |  |  |  |  |
| Ancient collection | 80<br>[sd=0] | 83.6<br>[sd=2.36] | 80<br>[sd=0] | 99.1<br>[sd=2.23] |
| Mixed collection | 80<br>[sd=0] | 83.6<br>[sd=2.25] | 80<br>[sd=0] | 103.2<br>[sd=2.47] |
| Recent collection | 80<br>[sd=0] | 83.5<br>[sd=2.38] | 80<br>[sd=0] | 108.3<br>[sd=2.36] |
| *Dynamic collections* |  |  |  |  |
| **Mobile recent** | 171.70<br>[sd=2.05] | 25.20<br>[sd=0.51] | 178.60<br>[sd=1.14] | 32.73<br>[sd=0.64] |
| **Mobile continuous** | 87.65<br>[sd=1.09] | 44.85<br>[sd=1.54] | 89.10<br>[sd=0.91] | 66.61<br>[sd=1.84] |
| **Mobile cumulative** | 199.80<br>[sd=3.04] | 44.01<br>[sd=1.10] | 215.05<br>[sd=1.61] | 47.71<br>[sd=0.62] |

Note: The random mating scenario is abbreviated with “*rm*” and the scenario which maximizes the genetic diversity is abbreviated with “*max\_GD*”.

**Figure 4.**
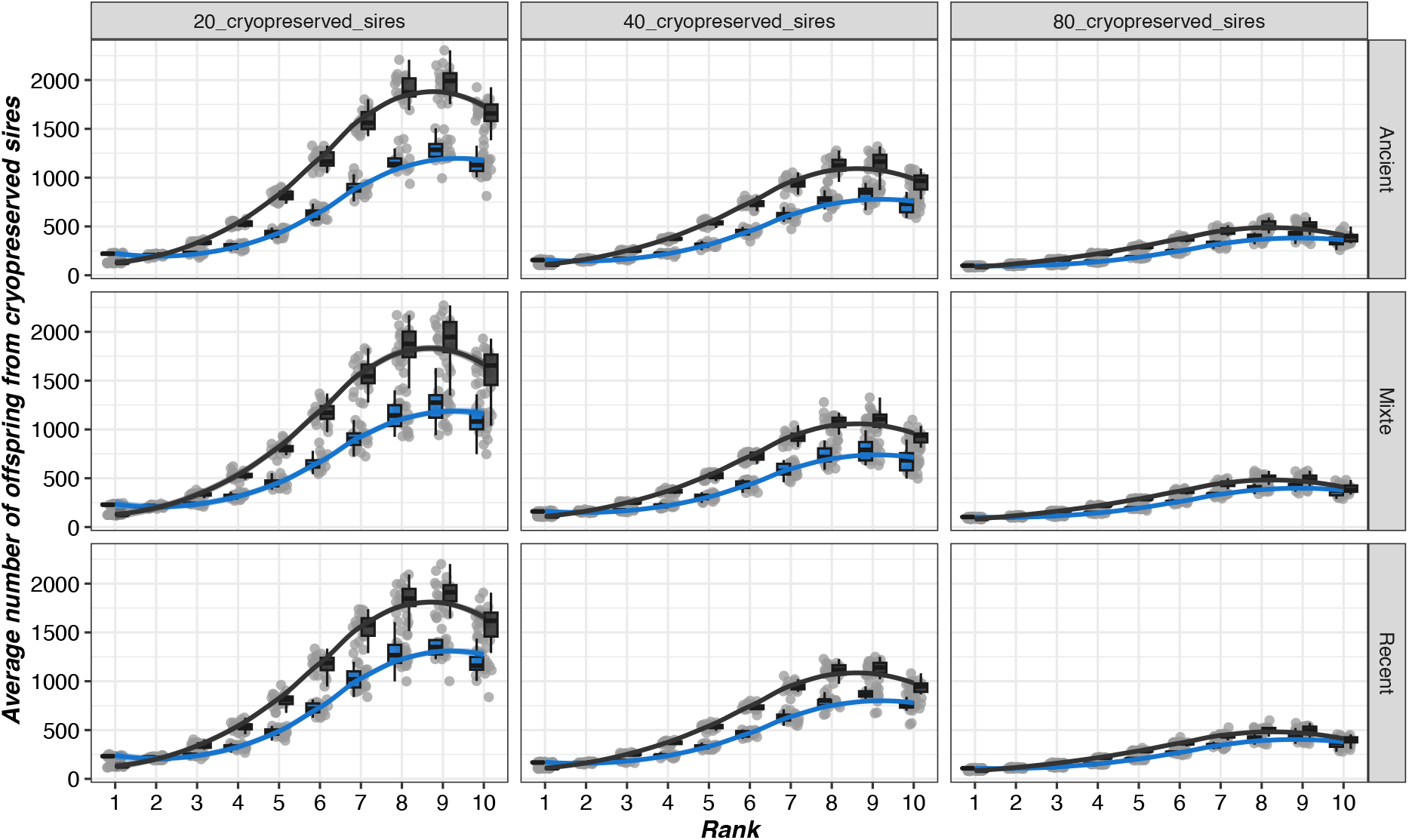
Number of offsprings of cryopreserved individuals across generations in a conservation program. The x-axis corresponds to the genealogical rank of individuals in relation to their cryopreserved sire. The random mating scenario (rm) is represented in black and the scenario which maximizes genetic diversity (max_GD) is represented in blue.

#### Impact on genetic diversity

In conservation programs which focusedg only on maintaining genetic diversity, there was a significant effect of the size of fixed collections (3-factor ANOVA test, F=659.7, df=346, p-val<0.05 for rm and F=229.5, df=346, p-val<0.05 for max_DG), in favor of larger collections (Figure 5A), while the age of collections had less impact on the trend in kinship (Figure 5B). However, when using dynamic collections, it is more advantageous to use older cryopreserved individuals (see Figure 5). The greatest impact on diversity was observed with the cumulative mobile collection (3-factor ANOVA test, F=410.6, df=230, p-val<0.05), which was the most extended in time and had the highest numbers of cryopreserved individuals.

**Figure 5.**
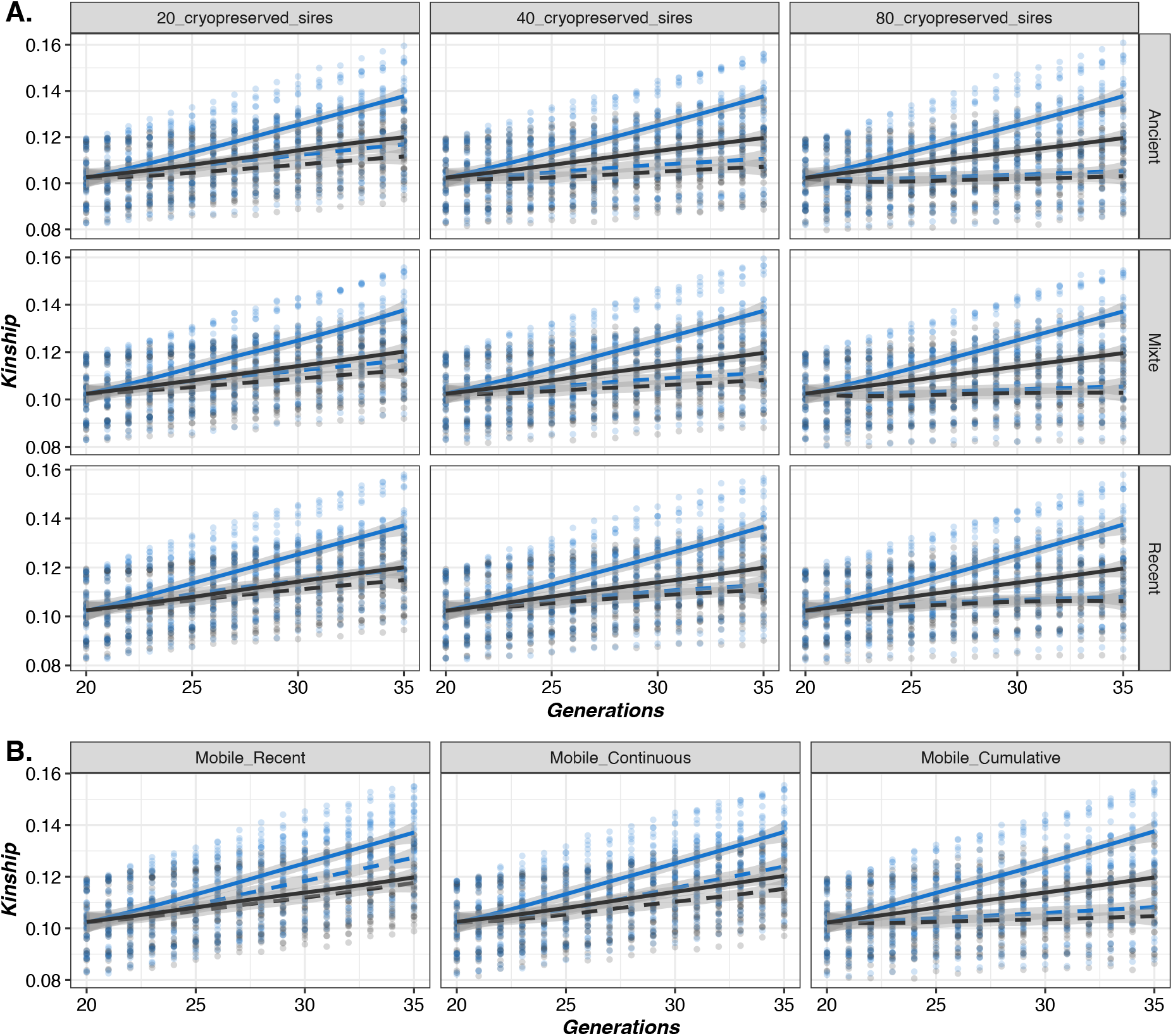
Evolution of average kinship over generations within populations under conservation according to the size (A) or the age of collections (B).The solid and dashed lines represent results without and with the use of *ex situ* collections, respectively. The random mating scenario (rm) is represented in black and the scenario which maximizes genetic diversity (max_GD) is represented in blue.

In both scenarios, (rm or max_DG), the genetic distance was consistently lower when using a cumulative mobile cryobank (between 2.14 × 10^−3^ and 2.23 × 10^−3^), whereas it was higher when using a recent or continuous mobile cryobank (between 3.87 × 10^−3^ and 5.52 × 10^−3^). Thus, genetic drift was globally more strongly reduced with the most complete collections (*i.e*. cumulative) than with the other two collections (2-factor ANOVA test, F=245.7, df=114, p-val<0.05 for rm and F=982.1, df=114, p-val<0.05 for max_GD), with lower Nei genetic distances between generations 20 and 35.

### Population under selection

Scenarios considered were those maximising breeding values, with optimal contribution selection (OCS) or without (max_BV).

#### Quantifying the use of ex situ resources

The most recent collections were used more, while the oldest collections were hardly used at all (Table 2 and Figure 6).The use of *ex situ* collections that spanned a long time interval was intermediate. Less markedly, the size of the cryopreserved collection had a positive impact on its use in breeding programs (3-factor ANOVA test, F=16.84, df=346, p-val<0.05). This trend was confirmed with the cumulative collection, which was the largest and most extended in time, and for which the cryopreserved individuals used were mainly the most recent (Figure 7). The other two mobile cryobanking strategies confirmed this trend, with recent mobile collections being much more widely used than continuous mobile collections (test ANOVA à 2 facteurs, F= 137.3, df=114, p-val<0.05), which included older generations but with a lower number of individuals cryopreserved per generation. However, it should be noted that, on average, few individuals were used in breeding programs, particularly when these did not incorporate constraints on genetic diversity. Nevertheless, the use of dynamic collections drastically increased their utilization as compared to fixed collections (Table 2).

**Table 2.** Number of cryopreserved individuals used and direct offspring produced in populations under selection (OCS and max_BV).

| CRYOBANK TYPE | OCS |  | max_BV |  |
| --- | --- | --- | --- | --- |
|  | No. cryopreserved sires used | No. offspring per cryopreserved sire | No. cryopreserved sires used | No. offspring per cryopreserved sire |
| *Fixed collections* |  |  |  |  |
| **20 cryopreserved sires** |  |  |  |  |
| Ancient collection | 0<br>[sd=0] | NA | 0<br>[sd=0] | NA |
| Mixed collection | 0.05<br>[sd=0.22] | 20<br>[sd=NA] | 0<br>[sd=0] | NA |
| Recent collection | 0.35<br>[sd=0.49] | 32.86<br>[sd=25.61] | 0.10<br>[sd=0.31] | 114<br>[sd=36.8] |
| **40 cryopreserved sires** |  |  |  |  |
| Ancient collection | 0<br>[sd=0] | NA | 0<br>[sd=0] | NA |
| Mixed collection | 0.20<br>[sd=0.41] | 20<br>[sd=7.12] | 0<br>[sd=0] | NA |
| Recent collection | 0.9<br>[sd=1.37] | 25.12<br>[sd=11.29] | 0.15<br>[sd=0.37] | 107.3<br>[sd=12.86] |
| **80 cryopreserved sires** |  |  |  |  |
| Ancient collection | 0<br>[sd=0] | NA | 0<br>[sd=0] | NA |
| Mixed collection | 0.70<br>[sd=0.80] | 23.80<br>[sd=15.40] | 0<br>[sd=0] | NA |
| Recent collection | 1.80<br>[sd=1.36] | 24.51<br>[sd=9.26] | 0.45<br>[sd=0.83] | 98.3<br>[sd=8.82] |
| *Dynamic collections* |  |  |  |  |
| **Mobile recent** | 11.10<br>[sd=2.81] | 26.15<br>[sd=3.98] | 0.40<br>[sd=0.60] | 111.71<br>[sd=15.98] |
| **Mobile continuous** | 3.80<br>[sd=1.61] | 25.98<br>[sd=8.29] | 0.05<br>[sd=0.22] | 108<br>[sd=NA] |
| **Mobile cumulative** | 8.40<br>[sd=2.95] | 24.24<br>[sd=4.39] | 0.20<br>[sd=0.41] | 106<br>[sd=20.72] |

Note: The scenario which maximize genetic progress is abbreviated with "max\_BV" and the scenario which maximizes genetic progress under a constraint of genetic diversity is abbreviated with "OCS".

**Figure 6.**
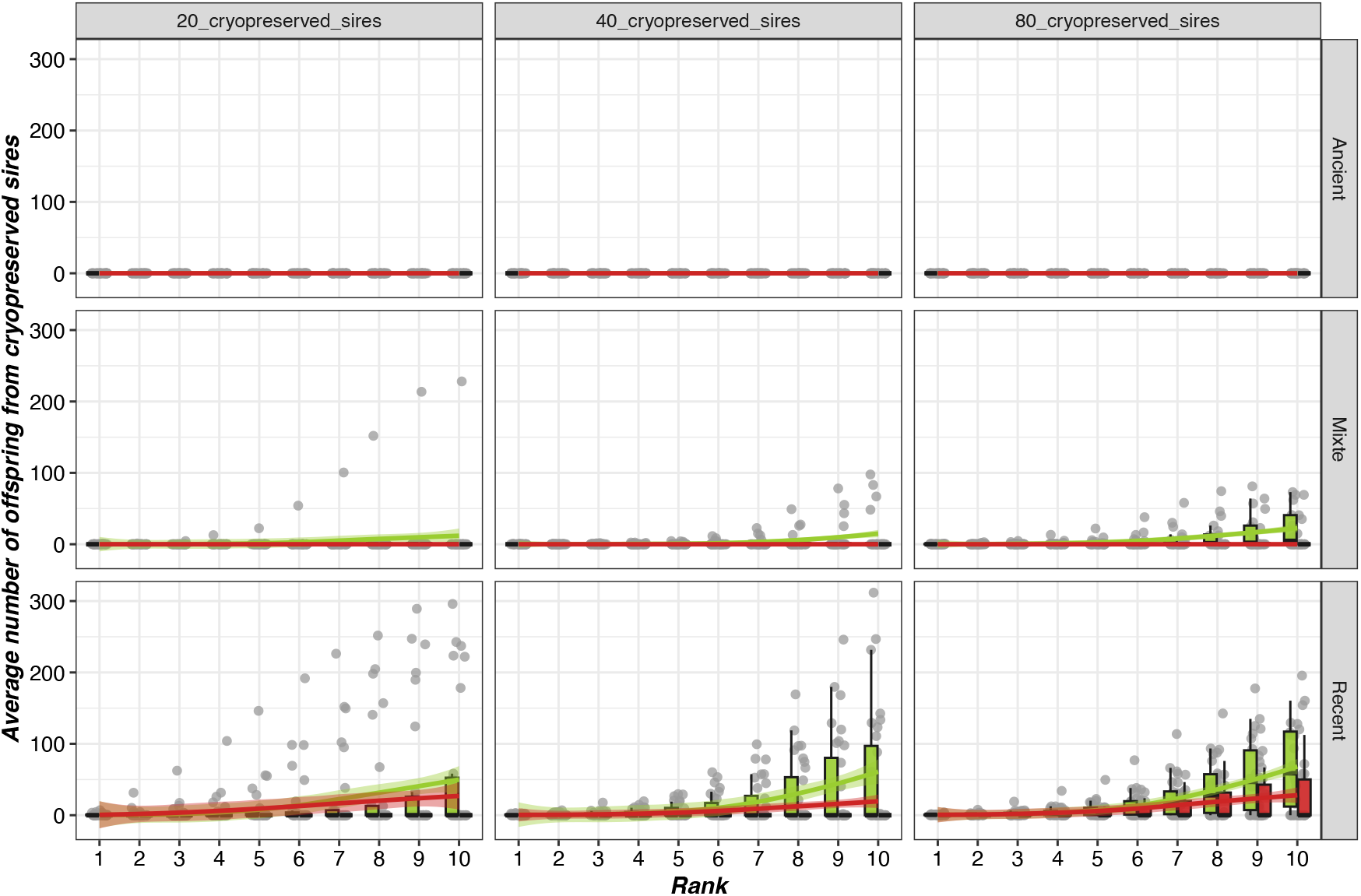
Number of cryopreserved individuals used and direct offspring produced in populations under selection (max_BV and OCS). The x-axis corresponds to the genealogical rank of individuals in relation to their cryopreserved ancestor from ex situ collections (i.e. rank 1 for children, rank 2 for grandchildren, rank 3 for great-grandchildren…).

**Figure 7.**
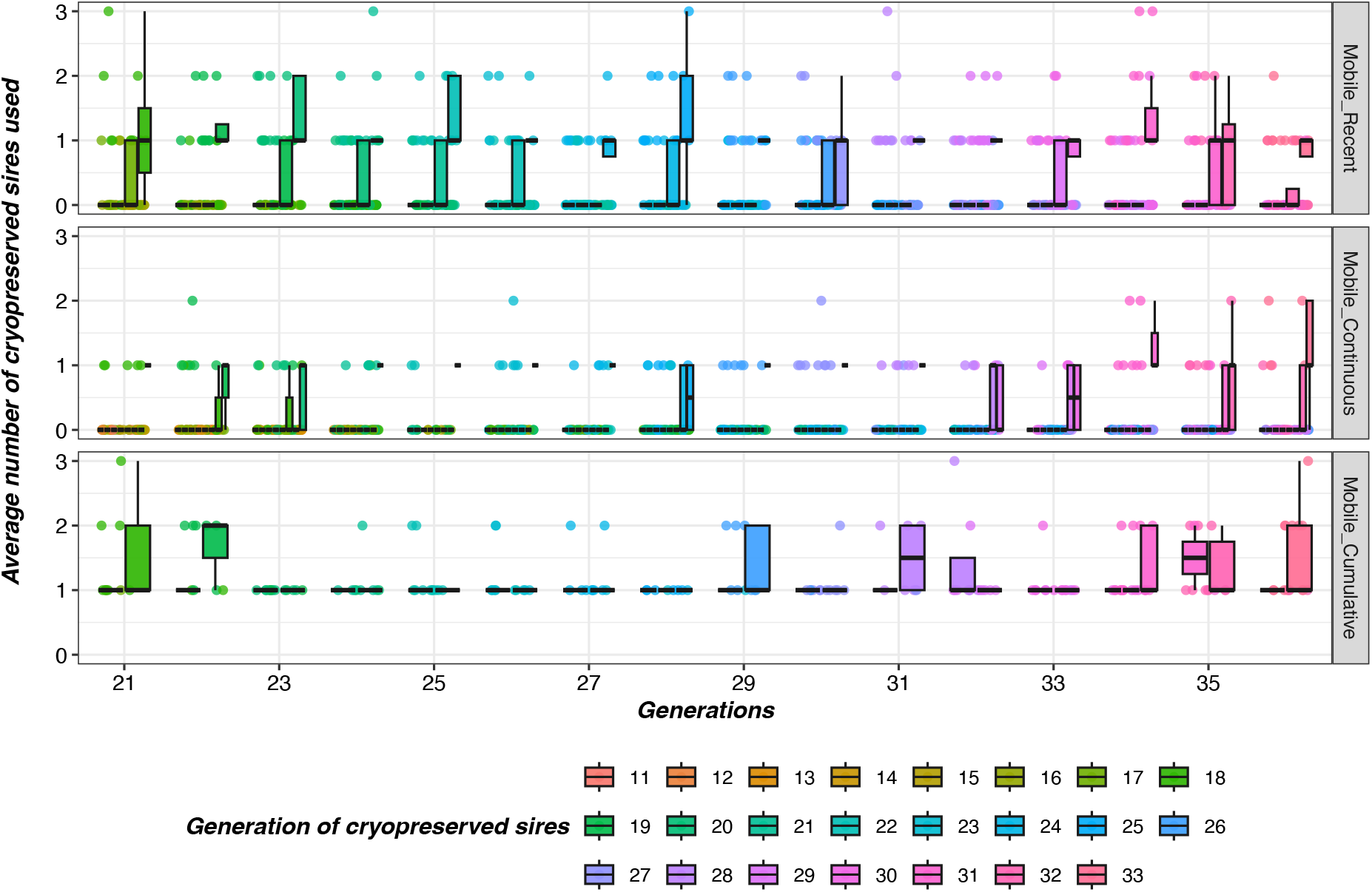
Age of genetic resources used in a program maximizing progress under diversity constraints (pop_OCS) with mobile cryobanks.

#### Impact on genetic diversity

For selection programs (max_BV and OCS), the age or size of fixed collections did not appear to have any effect on the evolution of genetic diversity within populations. However, when using dynamic collections, there was an advantage of cumulative strategies and recent collections, over those extended over time for the program aimed solely at improving performance, max_BV (see Figure 8). In the latter case, collections were used very little, which may explain this lower effect. In breeding programs involving genetic diversity (*i.e*. OCS), the impact of *ex situ* collections on genetic diversity was positive overall, although low, and did not differ between the different strategies (t-ratio=0.58, p-val=0.57).

**Figure 8.**
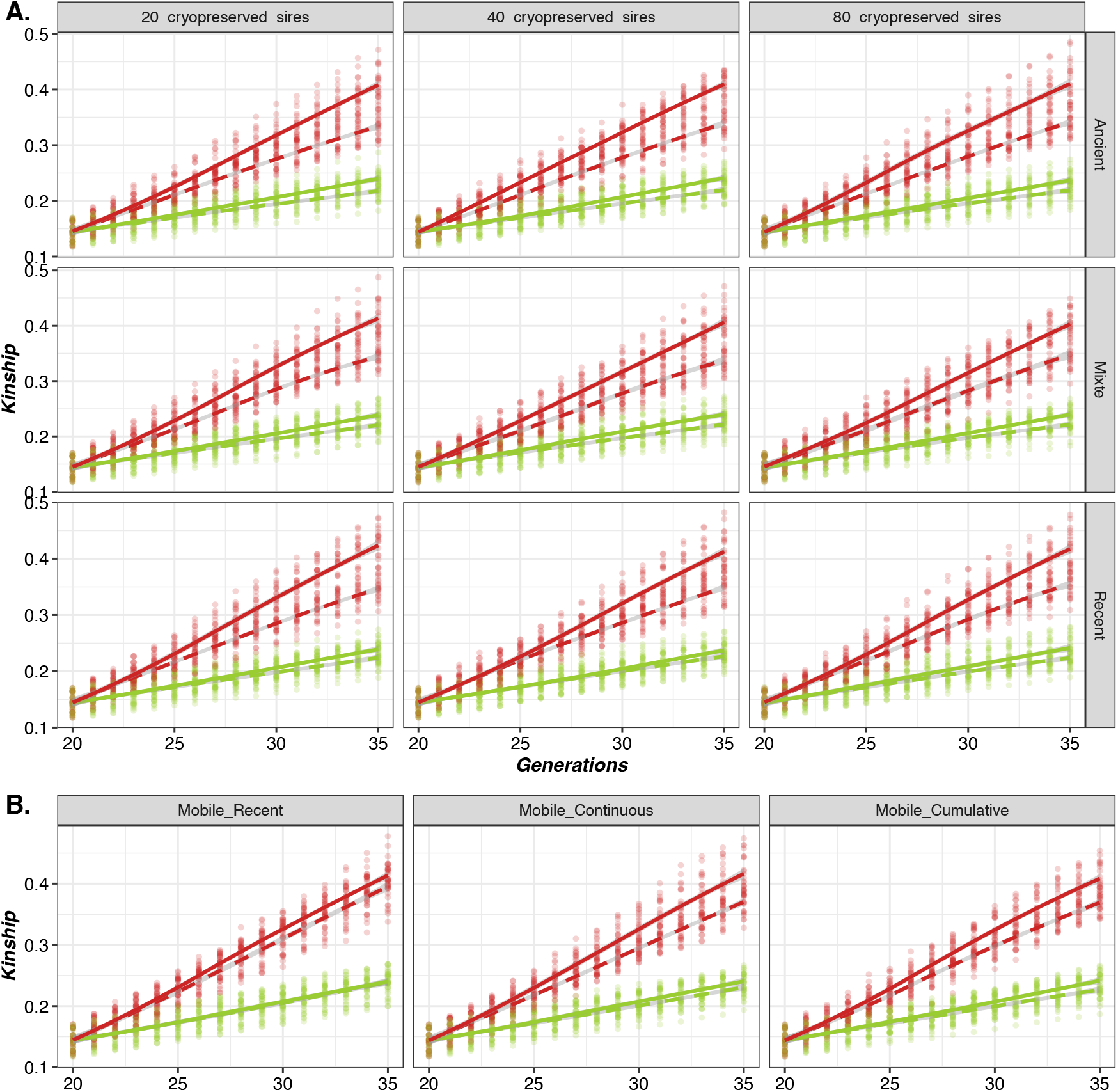
Evolution of average kinship over generations within selected populations according to breeding scenarios and constitution strategies of cryopreserved collections. The solid and dashed lines represent results without and with the use of *ex situ* collections, respectively. The scenario which maximizes genetic gain (max_BV) is represented in red and the scenario which maximizes genetic gain under a constraint of genetic diversity (OCS) is represented in green.

Finally, for a management program involving a constraint on diversity (*i.e*. OCS), genetic drift was globally more strongly reduced with the most complete collections (*i.e*. cumulative) than with recent collections (2-factor ANOVA test, F=22.84, df=114, p-val<0.05), with lower Nei genetic distances between generations 20 and 35 (see Table 3). On the other hand, for the program aimed solely at improving performance (*i.e*. max_BV), there were no differences between the different strategies to build collections.

**Table 3.** Genetic distances (Nei distance) in populations under selection between generations 20 and 35 according to the management program and the strategy used to constitute the cryobank.

| Cryobank types | pop_OCS | pop_max_BV |
| --- | --- | --- |
| Mobile recent | 2.00E-02<br>[sd=1.87E-03] | 5.86E-02<br>[sd=7.83E-03] |
| Mobile continuous | 1.72E-02<br>[sd=1.46E-03] | 5.23E-02<br>[sd=4.38E-03] |
| Mobile cumulative | 1.56E-02<br>[sd=1.09E-03] | 5.23E-02<br>[sd=4.76E-03] |

Note: The scenario which maximize genetic progress is abbreviated with “max\_BV” and the scenario which maximizes genetic progress under a constraint of genetic diversity is abbreviated with “OCS”.

#### Impact on genetic progress

The synthetic index was only slightly affected by the strategies used to build fixed cryopreserved collections, whatever their size or age (Figure 9) (3-factor ANOVA test, F=7.86, df=346, p-val<0.05 for max_BV and F=1.26, df=346, p-val=0.24 for OCS).

**Figure 9.**
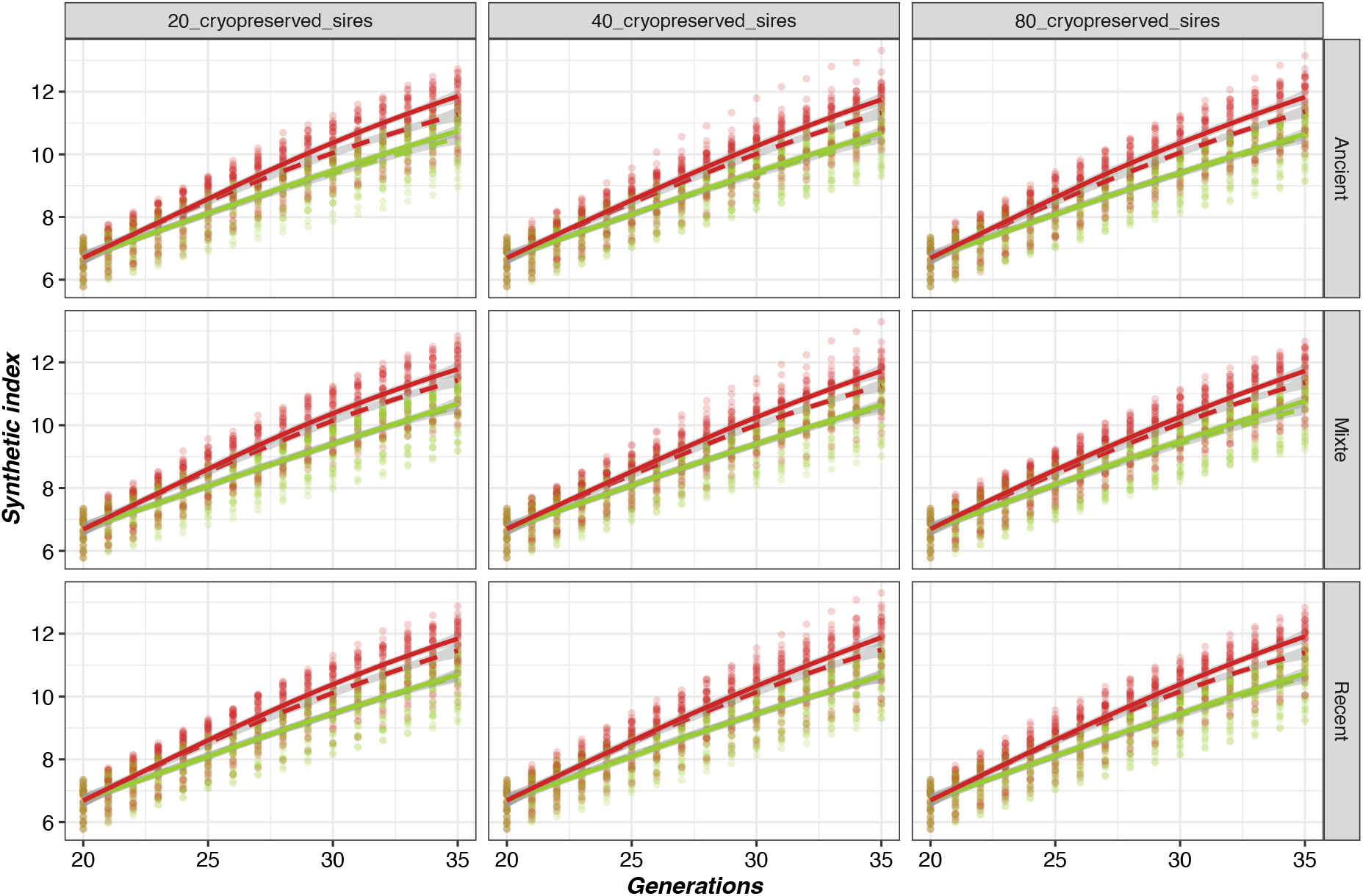
Evolution of the synthetic index value over generations according to the different strategies for building up fixed cryopreserved collections. The solid and dashed lines represent results without and with the use of *ex situ* collections, respectively. The scenario which maximizes genetic gain (max_BV) is represented in red and the scenario which maximizes genetic gain under a constraint of genetic diversity (OCS) is represented in green.

However, the impact on each trait differed (Figure 10, Figure 11). The use of cryopreserved individuals led to a slowdown in genetic progress for Trait 1 (3-factor ANOVA test, F=39.65, df=230, p-val<0.05) but an improvement in Trait 2 (3-factor ANOVA test, F=41.63, df=230, p-val<0.05), which was the minority trait in the index, negatively correlated with the majority trait. Thus, the larger the collection and the more ancient individuals it contains, the greater this impact will be.

**Figure 10.**
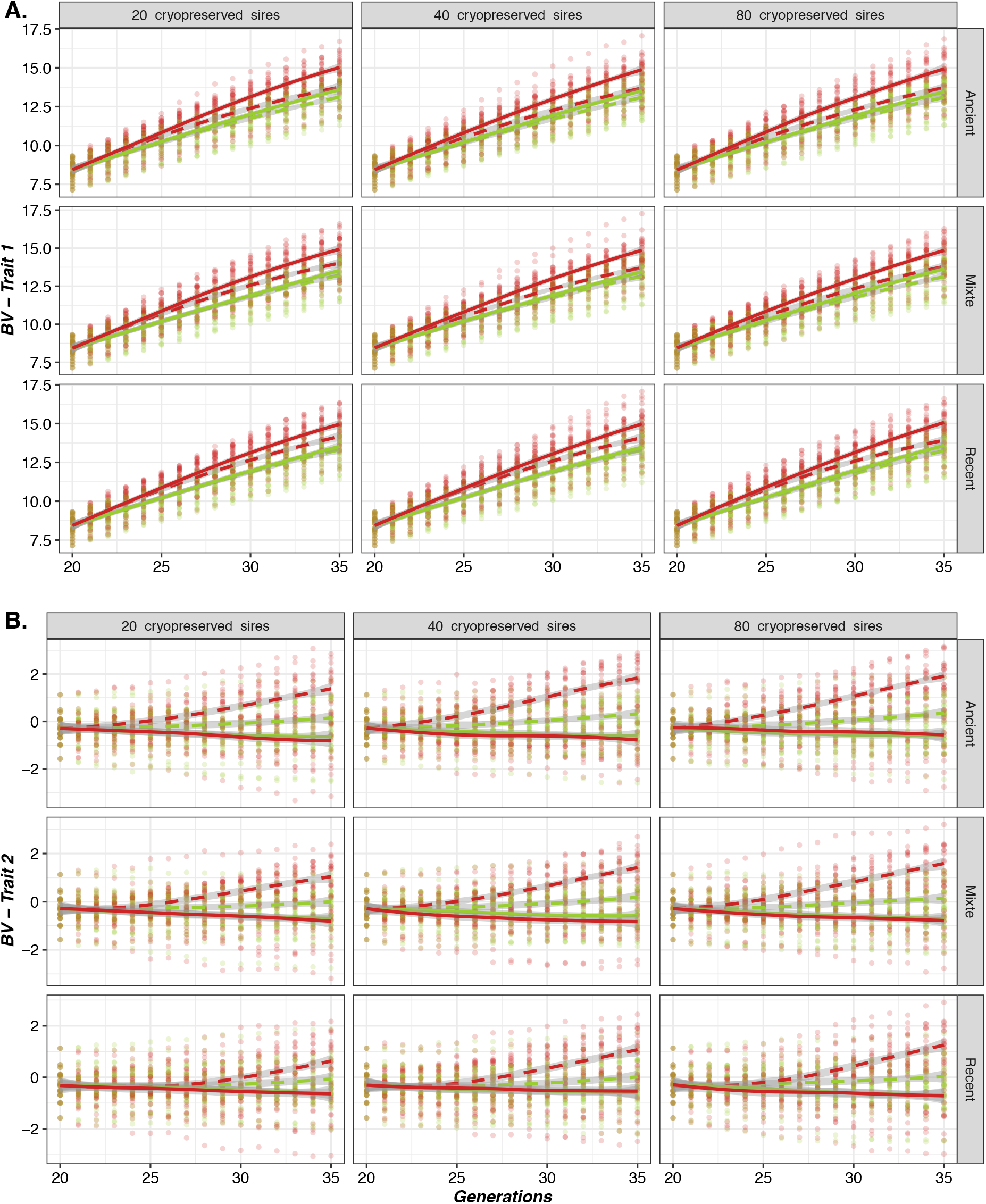
Evolution of the genetic value for each trait over generations according to different strategies for building fixed cryopreserved collections. The solid and dashed lines represent results without and with the use of *ex situ* collections, respectively. The scenario which maximizes genetic gain (max_BV) is represented in red and the scenario which maximizes genetic gain under a constraint of genetic diversity (OCS) is represented in green.

**Figure 11.**
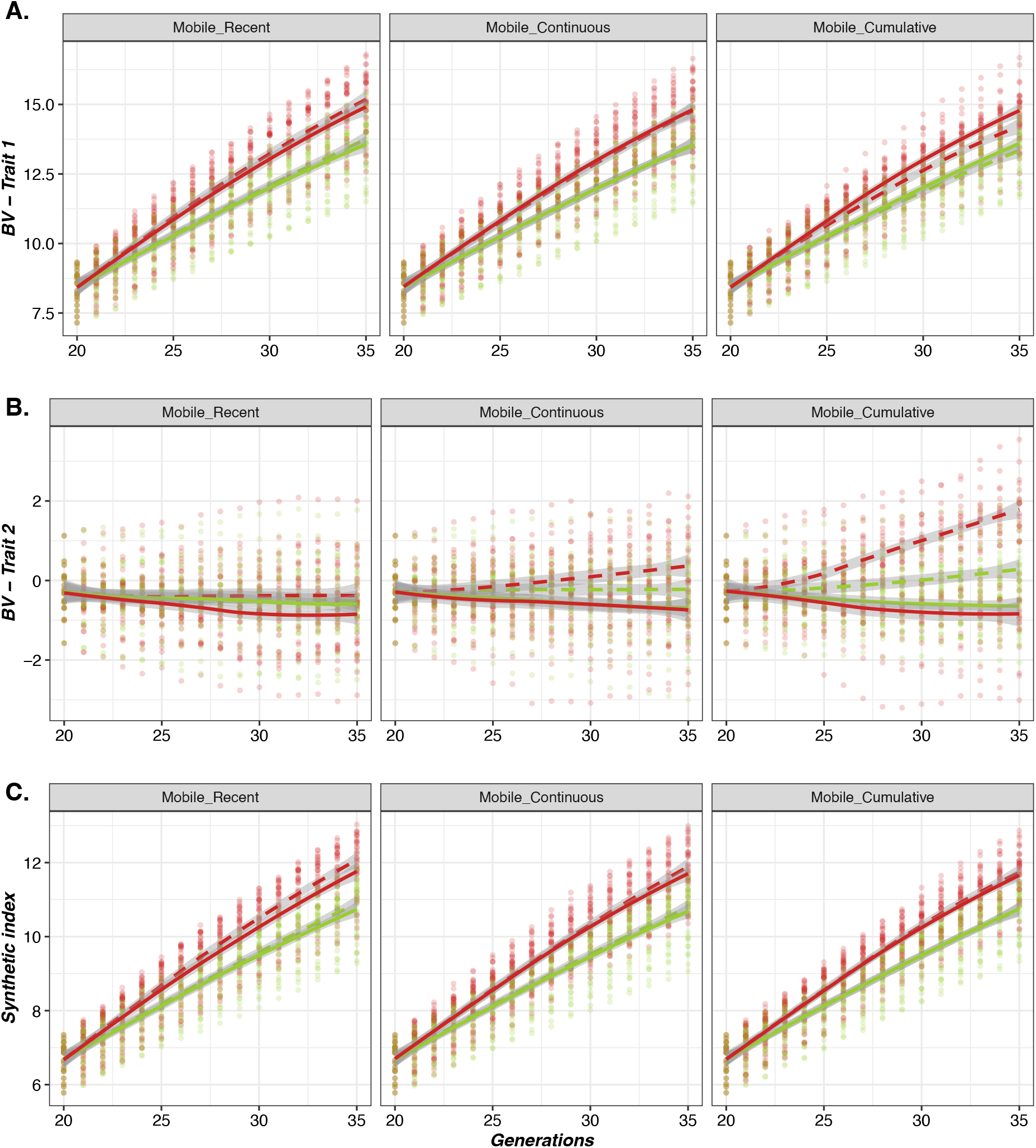
Evolution of the genetic value of the two traits over the generations according to the different strategies for building dynamic cryopreserved collections. The solid and dashed lines represent results without and with the use of *ex situ* collections, respectively. The scenario which maximizes genetic gain (max_BV) is represented in red and the scenario which maximizes genetic gain under a constraint of genetic diversity (OCS) is represented in green.

#### Impact on genetic variance

For fixed cryobanks, there were no significant differences in the trend of genetic variances for Trait 1 and Trait 2 between generations 20 and 35 with scenarios using OCS (3-factor ANOVA test, F=1.09, df=346, 16 p-val=0.37 for Trait 1 and F=1.19, df=346, p-val=0.19 for Trait 2). Details of the values for the various metrics are given in Additional file 1.

For mobile collections, the additive genetic variances of Trait1 and Trait2 were not significantly different according to the use of cryopreserved individuals for the three types of germplasm mobile collections in the OCS scenarios (2-factor ANOVA test, F=0.42, df=114, p-val=0.83 for Trait 1 and F=1.29, df=114, p-val=0.28 for Trait2) and in the max_BV scenarios (2-factor ANOVA test, F=1.98, df=114, p-val=0.09 for Trait 1 and F=0.18, df=114, p-val=0.97 for Trait2) (see Figure 12).

**Figure 12.**
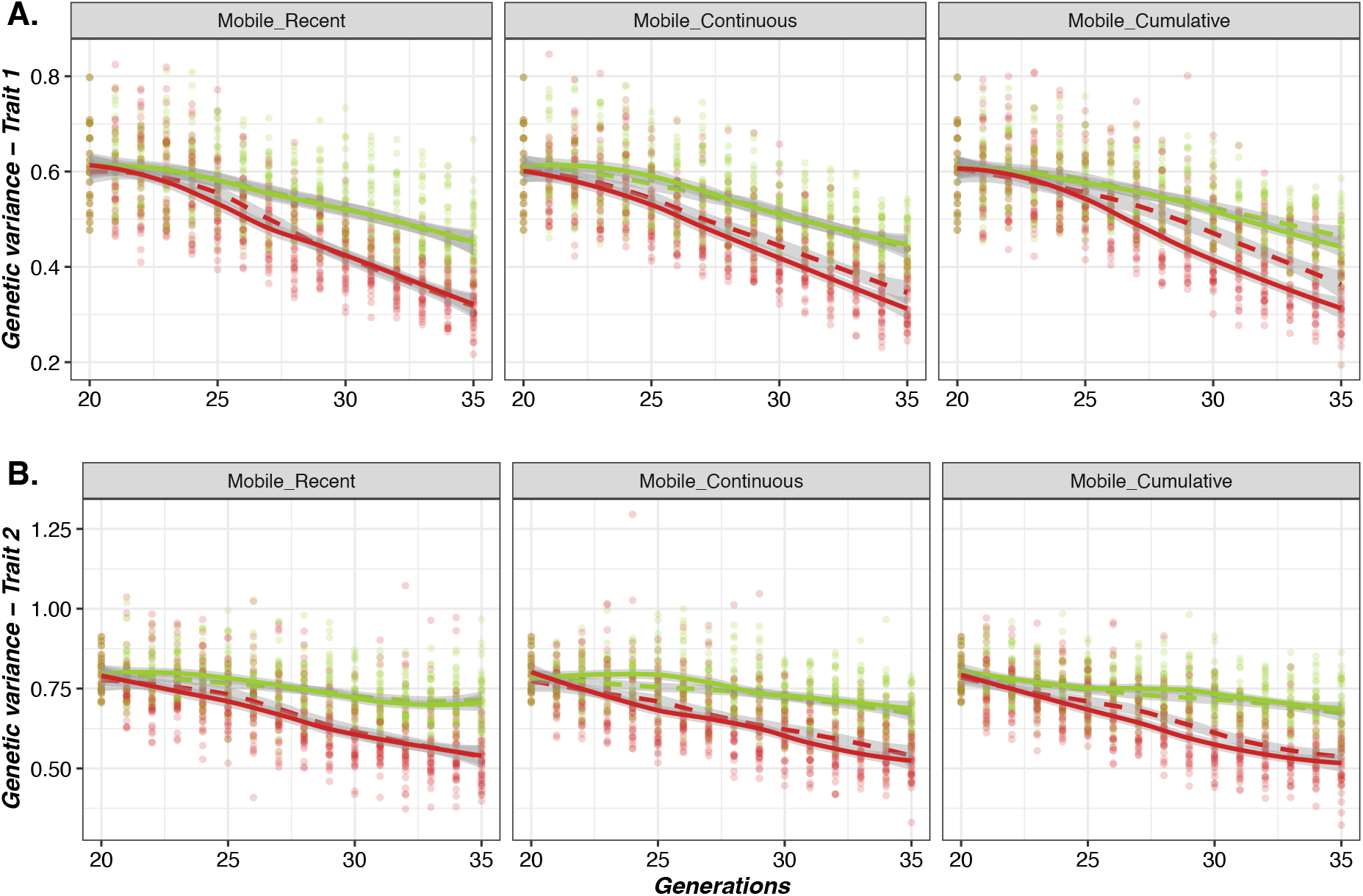
Evolution of genetic variances for trait 1 (A) and Trait 2 (B) according to the type of mobile cryobank for selected populations.

The solid and dashed lines represent results without and with the use of *ex situ* collections, respectively. The scenario which maximize genetic gain (max_BV) is represented in red and the scenario which maximizes genetic gain under a constraint of genetic diverisy (OCS) is represented in green.

## Discussion

### Collection strategy for a breed under selection

For populations under selection, the age of cryopreserved individuals seems to be more important than the size of *ex situ* collections. Indeed, genetic values of old cryopreserved individuals remain far behind those of contemporary candidates which greatly penalizes genetic progress, whereas for *ex situ* collections containing recent individuals, this performance gap is more moderate. When improving genetic progress is the main objective, a recent cryobank with many individuals maximizes the chances of having cryopreserved individuals with interesting genetic values. In this situation, old cryopreserved genetic resources are not relevant for immediate use, but remain very interesting for other purposes such as reorientation of the breeding scheme as previously discussed (Jacques et al., 2026) or reintroduction of lost lineages (Jacques et al., 2023). Consequently, for selected populations, older genetic resources may become relevant, as Leroy et al. (2011) pointed out, when there is a change in breeding objectives or a need to limit genetic drift, best achieved by the mobile cumulative gene bank.. In the case of fixed storage capacity, optimization of collections may require discarding some but not all old samples, using the Index of Diversity Impact (Jacques et al., 2024) to priorize the sires to be conserved. Such an optimization requires the gene bank to maintain regular contacts with the breeding scheme in order to anticipate its needs.

### Collection strategy for a breed in conservation

The use of cryopreserved resources is a real advantage for conservation programs, whatever the composition of the cryobank. The use of old cryopreserved individuals not only allows diversity to be reintroduced, but helps to maintain a correct level of genetic variability, since genetic gain is not a priority. This differs greatly from selected populations, and leads to a very interesting added value of old *ex situ* genetic resources, since the problem of lagging performance no longer arises. The regular use of cryopreserved genetic resources slows down the increase in inbreeding and the effect of genetic drift, all the more so when cryopreserved collections contain a large choice of individuals, as this allows better mixing of matings at each generation, drastically increasing the number of available breeding sires. The use of a cryobank containing few individuals is less efficient to slowdown the increase of inbreeding. The use of *ex-situ* genetic resources is therefore relevant for the conservation of endangered breeds, but the optimal collection-building strategy depends mainly on the number of donors available for conservation. A cryobank with a large number of aged individuals enables better management of matings and avoids too rapid an increase in inbreeding over the long term, while the use of a more moderate number of aged individuals offers fewer possibilities for managing genetic diversity. Thus, regular sampling of semen donors in each sire family should be recommanded to represent breed variability and increase the choice of sires for reproduction at each generation.

### The importance of economic and technical factors in the constitution of cryopreserved collections

For all scenarios, a few cryopreserved individuals were used in each generation, demonstrating that effective long-term management is only possible with a planned use of cryopreserved genetic resources over several generations. The cost of mobilizing *ex situ* collections should also be considered, particularly for some species or local breeds where logistics for using frozen semen is less widespread than for cattle. Consequently, a compromise might be to use a few individuals from the cryobank with contemporary breeders, then change donors at each generation. This system would be similar to the rotational mating strategy (Windig and Kaal, 2008; Theodorou and Couvet, 2010). This possibility could be simulated in order to study its impact, in particular with constraints in terms of the quantity of material (*e.g*. AI doses) to be used for each breeder, so as not to deplete stocks too quickly. Indeed, collections of certain breeds may have been difficult to build up. In the case of the French populations, some breeds have benefited from financial support from the CRB-Anim project to build up or enrich their collections, which shows that stocking strategies must also take into account the heterogeneous financial resources of different species and breeds. In this context, it may be appropriate to implement a dynamic management of the gene bank, balancing a regular use of collections with the preservation of old resources, in order to keep options for the future..

### Integrated strategy

A gene bank must address the needs of all population types. In case of discontinuous collection, we have seen that best sampling strategy will depend on the status of populations, either under selection or under conservation. The cumulative cryobank, which consists in continuously enriching collections, seems to be the best strategy as it combines large size (*i.e*. genetic variance), originality via older individuals (*e.g*. beneficial for new or minority traits) and good connection with current populations via recent individuals (*i.e*. limited performance gap). It is suitable for all types of population management programs, whether aimed at conservation or selection, with or without changes in objectives. This is currently the case for the French National Cryobank, which is therefore of undoubted interest for the future of animal populations, particularly in the context of the agroecological transition. A comprehensive sampling strategy may also better address the objective of restoration of a lost breed. However, this strategy is the most costly to set up and may not be sustainable on the long term where storage capacity may become limited. It is therefore necessary to consider more parsimonious scenarios, while maximizing their usefulness in the management of animal populations.

For selected populations, where the main objective is genetic progress, cryobanks are rarely used. It is mainly when selection objectives change that cryopreserved collections become most useful. In this case, preserving reproductive material from the last few generations prior to the current generation would be the most appropriate storage strategy (*i.e*. little gap in genetic values). This could avoid too rapid an increase in inbreeding over time, while retaining potential for new traits introduced for the new purpose. This management strategy is already known for private collections in cattle breeding companies, where old stocks are regularly replaced by contemporary bull semen. Thus, storage strategies could be complementary between public and private players. We could envisage extensive dynamic collections, but with variable numbers of individuals, depending on their age. In this way, a dynamic collection that samples a wide range of generations upstream, but with more recent individuals and fewer older ones, could be an optimal solution and should be tested at a later date. Decreasing the number of reproductive doses collected per sire could also be considered in order to decrease storage costs, since the number of direct progeny (rank 1) from cryopreserved bulls was generally small, though this was true only for conservation populations. In our study, donors also had an unlimited stock of semen doses; this factor should be taken into account, as it could influence the use of particular donors if the material stored in the collection is insufficient.

### Perspetives

It would be interesting to use simulations to explore more heterogeneous situations in terms of the type of individuals cryopreserved. Indeed, the *ex situ* genetic resources in our simulations were a sample of breeding males from some generations, meaning that there was a good chance that they were connected to the contemporary population. This situation corresponds to type III as defined by the French National Cryobank (Danchin Burge et al., 2006), where the aim is to have a representation of the breed’s diversity over a given period. Type II, created for entries of original individuals (*e.g*. family origins, particular genotypes, breeding values etc.) has not been considered, but could be of interest from the point of view of reorienting selection objectives. For example, it would be possible to stratify the sampling of individuals by selecting a few individuals whose genetic values show a large deviation from their generation average, or by taking into account new original traits.

Also, we considered here semen collections only, because this is the most frequent biological material currently stored, whereas embryos may be collected, at least in some species. Embryos are particularly relevant for the reconstitution of lost breeds, which can be at achieved faster than with semen only, where 4 generations are needed to restore more than 93% of the genome. Still, semen may be more suited to the real time management of diversity, particularly for local breeds.

## Conclusions

In conclusion, strategies for the collections management of *ex situ* animal genetic resources have to adjusted to the breeding program. For populations in selection, based only on the criterion of managing genetic progress and genetic diversity, keeping very old semen samples should be minimized, given that there is very little chance of these resources being used. For populations in conservation, the number of donors in collections greatly impact the potential for reintroducing genetic diversity. The regular conservation of representative sires at each generation would enable a wider range of uses, whatever the type of breeding scheme. Continuous enrichment of collections seems to be the best strategy for all breeds, provided that the relevance of old collections is regularly assessed, in order to control the total volume of samples in the gene bank.

## Acknowledgements

This study was partially funded by INRAE and the project GenResBridge. GenResBridge has received funding from the European Union’s Horizon 2020 research and innovation programme under grant agreement no. 817580. The PhD of AJ was funded by IDELE (Institut de l’Élevage), IFIP (Institut Français du Porc), SCC (Société Centrale Canine), and GenResBridge.

We are grateful to the INRAE MIGALE bioinformatics facility (MIGALE, INRAE, 2020. Migale bioinformatics Facility, doi: 10.15454/1.5572390655343293E12) for providing help and computing resources. The authors would like to acknowledge Torsten Pook for his assistance with the MoBPS software.

## Conflict of interest statement

The authors declare that they have no conflicts of interest.

## Notes

### Competing Interest Statement

The authors have declared no competing interest.

### Summary of Updates

The article has been revised to improve the clarity of the text and explanations, and the references have been updated.

