## Additional file 1 for "Managing the genetic diversity of animal populations using cryobanks: optimizing the constitution of *ex situ* collection?"

**Table S1:** Statistical metrics for each type of germplam collection for max_BV scenarios in a selected population.

| **CRYOBANK TYPE** | **Without use of cryopreserved genetic resources** | | | | | | | **With use of cryopreserved genetic resources** | | | | | | |
| --- | --- | --- | --- | --- | --- | --- | --- | --- | --- | --- | --- | --- | --- | --- |
|  | **∆BV1** | **∆BV2** | **∆Index** | **∆var_BV1** | **∆var_BV2** | **∆K** | **Genetic distance** | **∆BV1** | **∆BV2** | **∆Index** | **∆var_BV1** | **∆var_BV2** | **∆K** | **Genetic distance** |
| **20 cryopreserved sires** |  |  |  |  |  |  |  |  |  |  |  |  |  |  |
| Ancient collection | 6.58E+00 | -5.43E-01 | 5.15E+00 | -2.85E-01 | -2.35E-01 | 2.64E-01 | 6.32E-02 | 5.32E+00 | 1.63E+00 | 4.59E+00 | -1.99E-01 | -1.99E-01 | 1.89E-01 | 4.29E-02 |
| *sd* | *5.76E-01* | *7.02E-01* | *3.97E-01* | *7.90E-02* | *1.27E-01* | *2.67E-02* | *7.88E-03* | *5.26E-01* | *5.84E-01* | *4.32E-01* | *7.73E-02* | *1.03E-01* | *1.75E-02* | *4.58E-03* |
| Mixte collection | 6.50E+00 | -5.41E-01 | 5.09E+00 | -2.76E-01 | -2.96E-01 | 2.69E-01 | 6.56E-02 | 5.60E+00 | 1.32E+00 | 4.75E+00 | -2.16E-01 | -2.50E-01 | 2.01E-01 | 4.52E-02 |
| *sd* | *7.24E-01* | *7.52E-01* | *4.73E-01* | *5.89E-02* | *7.44E-02* | *2.62E-02* | *7.76E-03* | *6.32E-01* | *6.11E-01* | *5.05E-01* | *8.02E-02* | *8.33E-02* | *1.45E-02* | *4.22E-03* |
| Recent collection | 6.52E+00 | -3.55E-01 | 5.14E+00 | -2.97E-01 | -2.79E-01 | 2.79E-01 | 6.68E-02 | 5.74E+00 | 8.71E-01 | 4.76E+00 | -2.22E-01 | -2.17E-01 | 2.03E-01 | 4.67E-02 |
| *sd* | *4.82E-01* | *8.82E-01* | *3.27E-01* | *5.38E-02* | *9.25E-02* | *2.82E-02* | *8.81E-03* | *5.50E-01* | *7.51E-01* | *4.30E-01* | *7.77E-02* | *6.50E-02* | *1.76E-02* | *3.85E-03* |
| **40 cryopreserved sires** |  |  |  |  |  |  |  |  |  |  |  |  |  |  |
| Ancient collection | 6.44E+00 | -4.87E-01 | 5.05E+00 | -2.78E-01 | -2.57E-01 | 2.64E-01 | 6.19E-02 | 5.26E+00 | 2.10E+00 | 4.63E+00 | -2.06E-01 | -2.31E-01 | 1.95E-01 | 4.46E-02 |
| *sd* | *6.20E-01* | *8.19E-01* | *4.46E-01* | *6.73E-02* | *9.39E-02* | *1.80E-02* | *4.64E-03* | *4.88E-01* | *5.21E-01* | *3.44E-01* | *7.13E-02* | *7.95E-02* | *2.10E-02* | *6.00E-03* |
| Mixte collection | 6.44E+00 | -5.54E-01 | 5.04E+00 | -2.83E-01 | -2.76E-01 | 2.62E-01 | 6.35E-02 | 5.29E+00 | 1.67E+00 | 4.57E+00 | -2.03E-01 | -2.39E-01 | 1.93E-01 | 4.40E-02 |
| *sd* | *5.74E-01* | *5.26E-01* | *4.55E-01* | *6.21E-02* | *1.18E-01* | *2.27E-02* | *6.23E-03* | *4.37E-01* | *3.64E-01* | *3.71E-01* | *7.14E-02* | *7.80E-02* | *1.58E-02* | *4.22E-03* |
| Recent collection | 6.54E+00 | -2.76E-01 | 5.18E+00 | -2.71E-01 | -2.57E-01 | 2.68E-01 | 6.63E-02 | 5.66E+00 | 1.33E+00 | 4.79E+00 | -2.43E-01 | -2.09E-01 | 2.05E-01 | 4.64E-02 |
| *sd* | *8.22E-01* | *6.34E-01* | *5.92E-01* | *7.28E-02* | *7.46E-02* | *2.77E-02* | *7.92E-03* | *5.93E-01* | *5.53E-01* | *4.90E-01* | *6.51E-02* | *1.06E-01* | *2.26E-02* | *5.57E-03* |
| **80 cryopreserved sires** |  |  |  |  |  |  |  |  |  |  |  |  |  |  |
| Ancient collection | 6.49E+00 | -2.77E-01 | 5.14E+00 | -2.82E-01 | -2.82E-01 | 2.67E-01 | 6.45E-02 | 5.29E+00 | 2.18E+00 | 4.67E+00 | -1.89E-01 | -2.34E-01 | 1.97E-01 | 4.45E-02 |
| *sd* | *7.24E-01* | *8.80E-01* | *5.22E-01* | *7.15E-02* | *8.47E-02* | *3.28E-02* | *8.74E-03* | *5.06E-01* | *4.69E-01* | *3.76E-01* | *8.69E-02* | *7.88E-02* | *2.18E-02* | *7.07E-03* |
| Mixte collection | 6.43E+00 | -5.05E-01 | 5.04E+00 | -2.87E-01 | -2.75E-01 | 2.58E-01 | 6.21E-02 | 5.35E+00 | 1.84E+00 | 4.64E+00 | -2.10E-01 | -2.39E-01 | 2.04E-01 | 4.67E-02 |
| *sd* | *5.56E-01* | *6.87E-01* | *4.10E-01* | *6.97E-02* | *1.10E-01* | *2.49E-02* | *6.46E-03* | *4.60E-01* | *3.85E-01* | *3.36E-01* | *7.93E-02* | *7.60E-02* | *2.37E-02* | *5.68E-03* |
| Recent collection | 6.64E+00 | -4.18E-01 | 5.23E+00 | -2.97E-01 | -3.18E-01 | 2.74E-01 | 6.74E-02 | 5.50E+00 | 1.52E+00 | 4.70E+00 | -2.27E-01 | -2.37E-01 | 2.11E-01 | 4.85E-02 |
| *sd* | *5.73E-01* | *7.38E-01* | *3.82E-01* | *6.57E-02* | *8.44E-02* | *2.59E-02* | *6.97E-03* | *5.30E-01* | *5.23E-01* | *3.72E-01* | *6.80E-02* | *1.18E-01* | *1.95E-02* | *4.13E-03* |

Note: All cryobank types correspond to those developed in the Method section.

∆BV1: the delta between the mean value of Trait 1 true breeding values in generations 20 and 35

∆BV2: the delta between the mean value of Trait 2 true breeding values in generations 20 and 35

∆Index: the delta between the mean Synthetic Index value of the generation 20 and generation 35

∆var_BV1: the delta between the mean value of Trait 1 genetic variance in generations 20 and 35

∆var_BV2: the delta between the mean value of Trait 2 genetic variance in generations 20 and 35

∆K: the delta between kinship in generations 20 and 35

Genetic distance: the Nei’s genetic distance between the generations 20 and 35.

**Table S2:** Statistical metrics for each type of germplam collection for OCS scenarios in a selected population.

| **CRYOBANK TYPE** | **Without use of cryopreserved genetic resources** | | | | | | | **With use of cryopreserved genetic resources** | | | | | | |
| --- | --- | --- | --- | --- | --- | --- | --- | --- | --- | --- | --- | --- | --- | --- |
|  | **∆BV1** | **∆BV2** | **∆Index** | **∆var_BV1** | **∆var_BV2** | **∆K** | **Genetic distance** | **∆BV1** | **∆BV2** | **∆Index** | **∆var_BV1** | **∆var_BV2** | **∆K** | **Genetic distance** |
| **20 cryopreserved sires** |  |  |  |  |  |  |  |  |  |  |  |  |  |  |
| Ancient collection | 5.17E+00 | -3.90E-01 | 4.06E+00 | -1.54E-01 | -1.03E-01 | 9.53E-02 | 1.98E-02 | 4.69E+00 | 4.19E-01 | 3.83E+00 | -1.49E-01 | -8.88E-02 | 7.32E-02 | 1.39E-02 |
| *sd* | *4.43E-01* | *4.82E-01* | *3.34E-01* | *5.94E-02* | *9.71E-02* | *9.08E-03* | *1.79E-03* | *4.10E-01* | *3.48E-01* | *3.19E-01* | *4.68E-02* | *5.69E-02* | *6.02E-03* | *9.07E-04* |
| Mixte collection | 5.10E+00 | -4.20E-01 | 4.00E+00 | -1.47E-01 | -1.01E-01 | 9.47E-02 | 2.01E-02 | 4.80E+00 | 2.61E-01 | 3.89E+00 | -1.37E-01 | -1.14E-01 | 7.60E-02 | 1.43E-02 |
| *sd* | *4.80E-01* | *4.57E-01* | *3.44E-01* | *5.66E-02* | *6.52E-02* | *9.59E-03* | *2.48E-03* | *4.23E-01* | *3.38E-01* | *3.23E-01* | *5.05E-02* | *8.02E-02* | *5.14E-03* | *9.38E-04* |
| Recent collection | 5.10E+00 | -3.71E-01 | 4.00E+00 | -1.59E-01 | -1.21E-01 | 9.52E-02 | 1.98E-02 | 4.88E+00 | 1.98E-01 | 3.94E+00 | -1.30E-01 | -1.05E-01 | 7.98E-02 | 1.51E-02 |
| *sd* | *4.73E-01* | *3.79E-01* | *3.48E-01* | *5.43E-02* | *7.94E-02* | *8.90E-03* | *2.19E-03* | *4.10E-01* | *3.76E-01* | *3.10E-01* | *5.85E-02* | *8.94E-02* | *6.20E-03* | *1.15E-03* |
| **40 cryopreserved sires** |  |  |  |  |  |  |  |  |  |  |  |  |  |  |
| Ancient collection | 5.09E+00 | -3.38E-01 | 4.00E+00 | -1.65E-01 | -8.48E-02 | 9.71E-02 | 2.05E-02 | 4.67E+00 | 5.80E-01 | 3.85E+00 | -1.30E-01 | -1.36E-01 | 7.49E-02 | 1.41E-02 |
| *sd* | *3.85E-01* | *5.08E-01* | *3.05E-01* | *6.48E-02* | *8.03E-02* | *1.10E-02* | *2.56E-03* | *4.13E-01* | *3.02E-01* | *3.39E-01* | *6.50E-02* | *6.82E-02* | *5.95E-03* | *1.01E-03* |
| Mixte collection | 5.01E+00 | -3.22E-01 | 3.95E+00 | -1.64E-01 | -1.02E-01 | 9.56E-02 | 2.00E-02 | 4.72E+00 | 4.46E-01 | 3.87E+00 | -1.34E-01 | -1.19E-01 | 7.71E-02 | 1.43E-02 |
| *sd* | *3.81E-01* | *3.74E-01* | *2.90E-01* | *6.09E-02* | *8.01E-02* | *7.72E-03* | *1.74E-03* | *4.48E-01* | *3.21E-01* | *3.69E-01* | *5.85E-02* | *7.09E-02* | *6.57E-03* | *1.11E-03* |
| Recent collection | 5.03E+00 | -3.06E-01 | 3.97E+00 | -1.56E-01 | -7.40E-02 | 9.29E-02 | 1.92E-02 | 4.85E+00 | 2.81E-01 | 3.94E+00 | -1.45E-01 | -1.20E-01 | 8.27E-02 | 1.58E-02 |
| *sd* | *4.77E-01* | *4.23E-01* | *3.57E-01* | *5.86E-02* | *9.73E-02* | *8.22E-03* | *1.72E-03* | *4.60E-01* | *3.29E-01* | *3.36E-01* | *5.24E-02* | *6.00E-02* | *7.08E-03* | *1.17E-03* |
| **80 cryopreserved sires** |  |  |  |  |  |  |  |  |  |  |  |  |  |  |
| Ancient collection | 5.02E+00 | -2.97E-01 | 3.96E+00 | -1.58E-01 | -1.34E-01 | 9.30E-02 | 1.92E-02 | 4.67E+00 | 6.15E-01 | 3.86E+00 | -1.21E-01 | -1.15E-01 | 7.52E-02 | 1.40E-02 |
| *sd* | *3.75E-01* | *4.18E-01* | *2.95E-01* | *5.49E-02* | *5.05E-02* | *8.45E-03* | *1.74E-03* | *4.09E-01* | *3.36E-01* | *3.10E-01* | *4.72E-02* | *6.24E-02* | *6.86E-03* | *1.19E-03* |
| Mixte collection | 5.21E+00 | -4.46E-01 | 4.08E+00 | -1.38E-01 | -1.06E-01 | 9.57E-02 | 1.98E-02 | 4.69E+00 | 4.16E-01 | 3.84E+00 | -1.42E-01 | -1.26E-01 | 7.72E-02 | 1.40E-02 |
| *sd* | *4.74E-01* | *4.95E-01* | *3.74E-01* | *5.41E-02* | *5.41E-02* | *8.19E-03* | *1.89E-03* | *4.50E-01* | *4.01E-01* | *3.53E-01* | *6.40E-02* | *6.51E-02* | *6.69E-03* | *1.27E-03* |
| Recent collection | 5.16E+00 | -4.40E-01 | 4.04E+00 | -1.57E-01 | -9.26E-02 | 9.76E-02 | 2.04E-02 | 4.81E+00 | 3.08E-01 | 3.91E+00 | -1.34E-01 | -1.40E-01 | 7.92E-02 | 1.52E-02 |
| *sd* | *4.62E-01* | *3.68E-01* | *3.51E-01* | *5.44E-02* | *8.31E-02* | *1.03E-02* | *2.17E-03* | *3.82E-01* | *3.89E-01* | *2.87E-01* | *5.94E-02* | *6.53E-02* | *7.35E-03* | *1.28E-03* |

Note: All cryobank types correspond to those developed in the Method section.

∆BV1: the delta between the mean value of Trait 1 true breeding values in generations 20 and 35

∆BV2: the delta between the mean value of Trait 2 true breeding values in generations 20 and 35

∆Index: the delta between the mean Synthetic Index value of the generation 20 and generation 35

∆var_BV1: the delta between the mean value of Trait 1 genetic variance in generations 20 and 35

∆var_BV2: the delta between the mean value of Trait 2 genetic variance in generations 20 and 35

∆K: the delta between kinship in generations 20 and 35

Genetic distance: the Nei’s genetic distance between the generations 20 and 35.
